# In silico modelling demonstrates that user variability during tumor measurement can affect in vivo therapeutic efficacy outcomes

**DOI:** 10.1101/2022.04.20.487864

**Authors:** Jake T. Murkin, Hope E. Amos, Daniel W. Brough, Karl D. Turley

## Abstract

User measurement bias during subcutaneous tumor measurement is a source of variation in preclinical in vivo studies. We investigated whether this user variability could impact efficacy study outcomes, in the form of the false negative result rate when comparing treated and control groups.

Two tumor measurement methods were compared; calipers which rely on manual measurement, and an automatic 3D and thermal imaging device. Tumor growth curve data were used to create an in silico efficacy study with control and treated groups. Before applying user variability, treatment group tumor volumes were statistically different to the control group. Utilizing data collected from 15 different users across 9 in vivo studies, user measurement variability was computed for both methods and simulation was used to investigate its impact on the in silico study outcome.

User variability produced a false negative result in 3.5% to 19.5% of simulated studies when using calipers, depending on treatment efficacy. When using an imaging device with lower user variability this was reduced to 0.0% to 2.4%, demonstrating that user variability impacts study outcomes and the ability to detect treatment effect.

Reducing variability in efficacy studies can increase confidence in efficacy study outcomes without altering group sizes. By using a measurement device with lower user variability, the chance of missing a therapeutic effect can be reduced and time and resources spent pursuing false results could be saved. This improvement in data quality is of particular interest in discovery and dosing studies, where being able to detect small differences between groups is crucial.

## Introduction

Subcutaneous tumor xenograft models are used to study tumor progression and responses in vivo. Tumor volume is the most commonly used metric to monitor progression of tumor and response to treatments. Tumor volume is calculated using two or more dimensions; most commonly length and width, using tumor width as a proxy for height^1^. These dimensions are measured manually (with calipers), or by using imaging techniques including MRI, CT, fluorescence, or 3D imaging combined with a thermal signature ^2–5^. Calipers are the most common tool of choice due to their low cost, however they produce less precise, more variable results than CT^3^, ultrasound^6^, and 3D and thermal imaging methods^5^ due to user measurement variation. Caliper users must determine the longest tumor length and its perpendicular width by eye which is highly subjective^7^, and the mechanical nature of calipers adds further variation by allowing the tumor to be squeezed, influencing its shape and recorded dimensions.

The user variability problem in tumor measurement has been addressed in the clinical field and has been found to affect MR and CT imaging techniques that rely on manual measurement methods^8,9^. Computer-aided tumor measurement can be used to more precisely assess tumor volume by removing user variability and bias, and software now exists to define and measure tumor dimensions automatically as part of imaging methods. Therefore, sources of user measurement variation can be removed by imaging methods that use algorithms and machine learning to determine the longest length and width automatically, and by designing tools that do not come into contact with the tumor. Partial or fully automatic image processing and tumor measurement has been widely adopted in oncology clinics, setting a precedent for this technology to improve data quality and throughput in preclinical trials^9,10^. Automatic tumor measurement with 3D and thermal tumor imaging has indeed been shown to significantly reduce user measurement variability in subcutaneous in vivo studies^5,11^.

Cancer research is affected by a reproducibility crisis, with estimates of published studies that can be reproduced by another team as low as 11%^12^. This irreproducibility stems from many sources including study design and data reporting^13,14^, variation within animal models^15^, and use of low quality or misidentified biospecimens and cell lines^16,17^. Lower precision during measurement also affects study reproducibility; caliper users cannot swap in and out of studies as the inter-operator variability is so high that measurements between users are often not comparable, even when measuring the same animal. Thus, reducing measurement variability is a promising option to achieve greater repeatability of results. Caliper measurement variability is a known problem, however their use is still ubiquitous, and the effects on study endpoints have not been investigated in detail.

High attrition of drugs during late-stage clinical trials is another prevalent problem in oncology, where only 5% of drugs in Phase I will be successfully licensed^18^. A greater focus of resources at the preclinical drug discovery stage (the ‘quick win, fast fail’ paradigm where drug candidates are filtered out during preclinical testing) has been suggested as a solution to reduce drug development costs^19^. More certainty of drug effects earlier on will reduce attrition and costs downstream, but this method is also dependent on a low false negative rate so that effective drugs are not discarded. Decreasing user variability is therefore a viable target to increase certainty in drug effects in the preclinical stage.

## Aims of the study

A common method used to evaluate treatment efficacy in subcutaneous tumor models is to compare average tumor volume of a treated group with that of a control (untreated) group on the final day of a study. Significant differences between groups are determined using a statistical test, for example a t-test. We hypothesized that larger user measurement variability would result in larger standard deviation of group volume and less consistency in group volumes when repeating a study and would therefore affect the conclusions made in the study and repeats.

As previously shown, user measurement variability and bias can affect preclinical in vivo tumor studies, from randomization to study outcomes ^5,11^. These variability data were used as a start point for this investigation in order to investigate how variability affects study endpoints in more detail. Mathematical modelling was chosen to create a controlled in vivo efficacy study scenario in which the only variation in study outcome was produced by the user variability applied to the model. In a typical in vivo study, other sources of variation including differences between individual rodents, and laboratory conditions could also affect the outcome, so this method allowed us to isolate the effects of user measurement variation on the study outcomes with confidence.

## Methods

To investigate the impact of inter-operator variability on study outcome and study reproducibility a combination of modelling and simulation was used to create an in silico study of tumor growth with two groups (control and treated). Study outcome was the detection of an effective treatment, defined as a statistical difference in group average tumor volumes on the final day of the study. An overview of the modelling and simulation process is shown in Figure 1.

**Figure 1:**
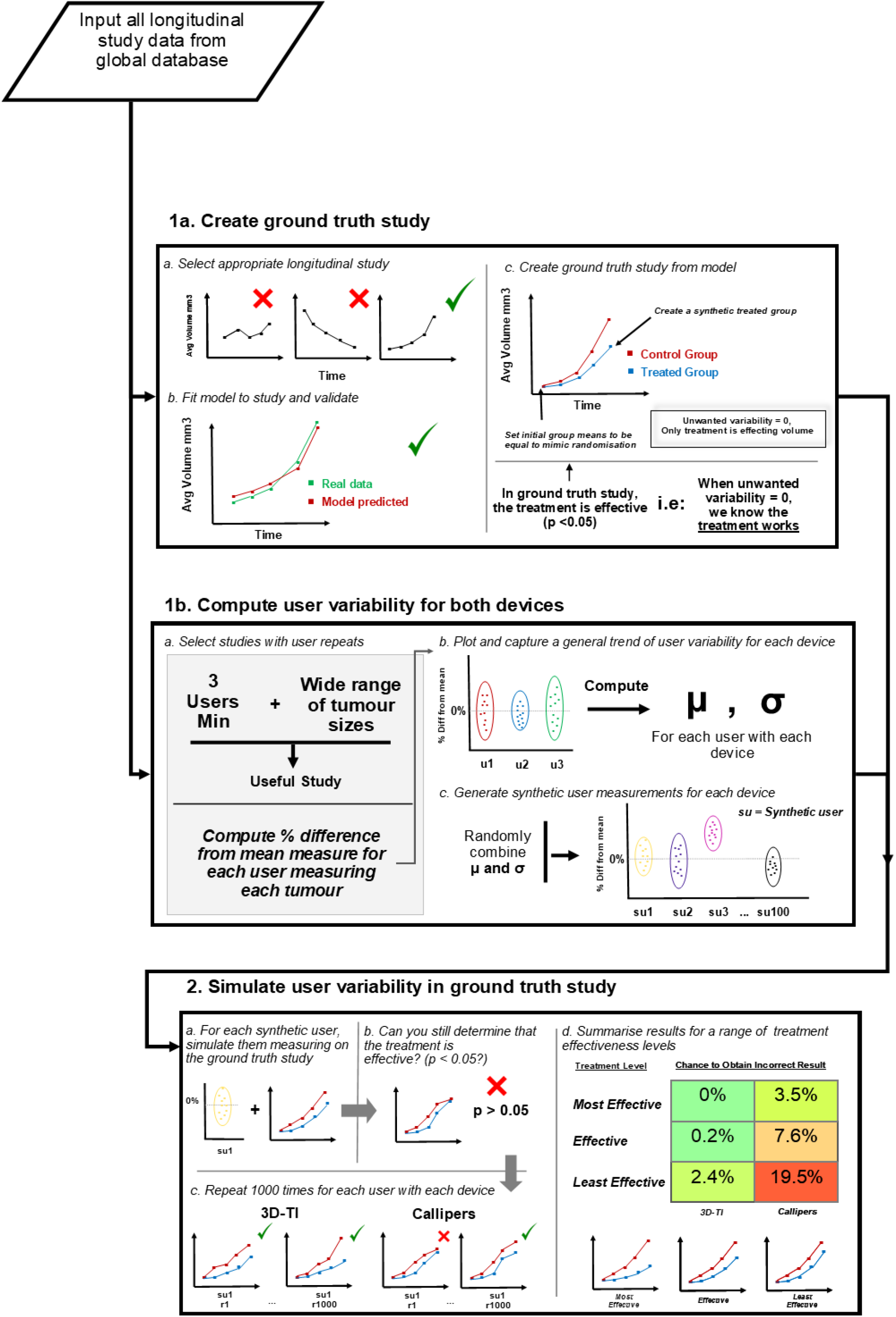
Flow chart describing the analysis process

### Tumor measurement devices

Calipers were used in the standard way; users chose the longest tumor length and its perpendicular width by eye, and measured along these axes by placing caliper blades around the tumor.

Stereoscopic RGB and thermal images were captured and converted into 3D tumor models using the BioVolume 3D and thermal imaging (3D-TI) device and software. BioVolume’s 3D-TI measurement algorithm determined the tumor’s length and width using the same automatic method every time, removing user bias. Details on measurement technique, the BioVolume system, and how scans were processed are available in our previous paper^11^.

Tumor volume was calculated from length and width in the same way for both devices, using the formula^20^:

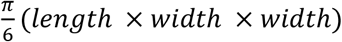

### In vivo data collection

The 3D-TI device and system (BioVolume) were used by 27 client organizations with training and support from Fuel3D. Studies were designed to compare variability of caliper measurements to the 3D-TI device. All animal care, lab work, caliper measurements and image scans were carried out by scientists in client organizations according to their own animal handling and ethics protocols. Data were shared with Fuel3D to use in an aggregated and anonymized way, forming the ‘global dataset’. Client companies and scientists did not have financial interests in BioVolume.

Appropriate in vivo longitudinal studies from this global dataset of 3D-TI measurements were selected as described in the following ‘ground truth’ modelling and ‘computing user variability’ sections.

## Creating the Ground Truth Study

### In vivo data processing

To create representative synthetic study data in which unwanted variability is equal to zero, a template growth curve for synthetic users to ‘measure’ was established (**Figure 1, Box 1a**). A tumor growth study where growth was measured using 3D-TI and calipers at 7 points across a 16-day period was used to model the template tumor growth curves. 3D-TI measurements had the lowest inter-operator variability (assessed by coefficient of variation, 0.181 vs 0.238, p=0.026, Wilcoxon – signed rank test) so were used for modelling. Repeat measurements enabled estimation of stable means when fitting the model.

### Fitting the model

Modelling synthetic data allowed removal of unwanted sources of variability. Any difference in average treatment between groups could then be confirmed to be from treatment.

A generalized linear model was fit to the study data in R. The model consisted of:

- A fixed group slope by day to estimate the growth rate of the group
- A fixed rodent intercept, to account for varying initial rodent volumes.

Combining the above gives us the model formula:

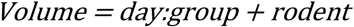

where *day* represents the day since first measurement. A generalized linear model was fit using a log link function and a gamma family to account for exponential tumor growth, and heteroskedastic growth within groups. The growth rate obtained from the *day:group* term in the model was 0.236 log units, or a daily increase in growth rate of exp(0.236) = 1.26.

To validate the model and confirm that the group growth rate obtained by the model accurately represented the in vivo study data, growth curves were plotted for both the in vivo study, and modelled study (**Figure 2a**). The growth curves were closely aligned, with overlapping CIs, indicating that the model represented the study well, confirming that the growth rate of the control group was captured successfully.

**Figure 2.**
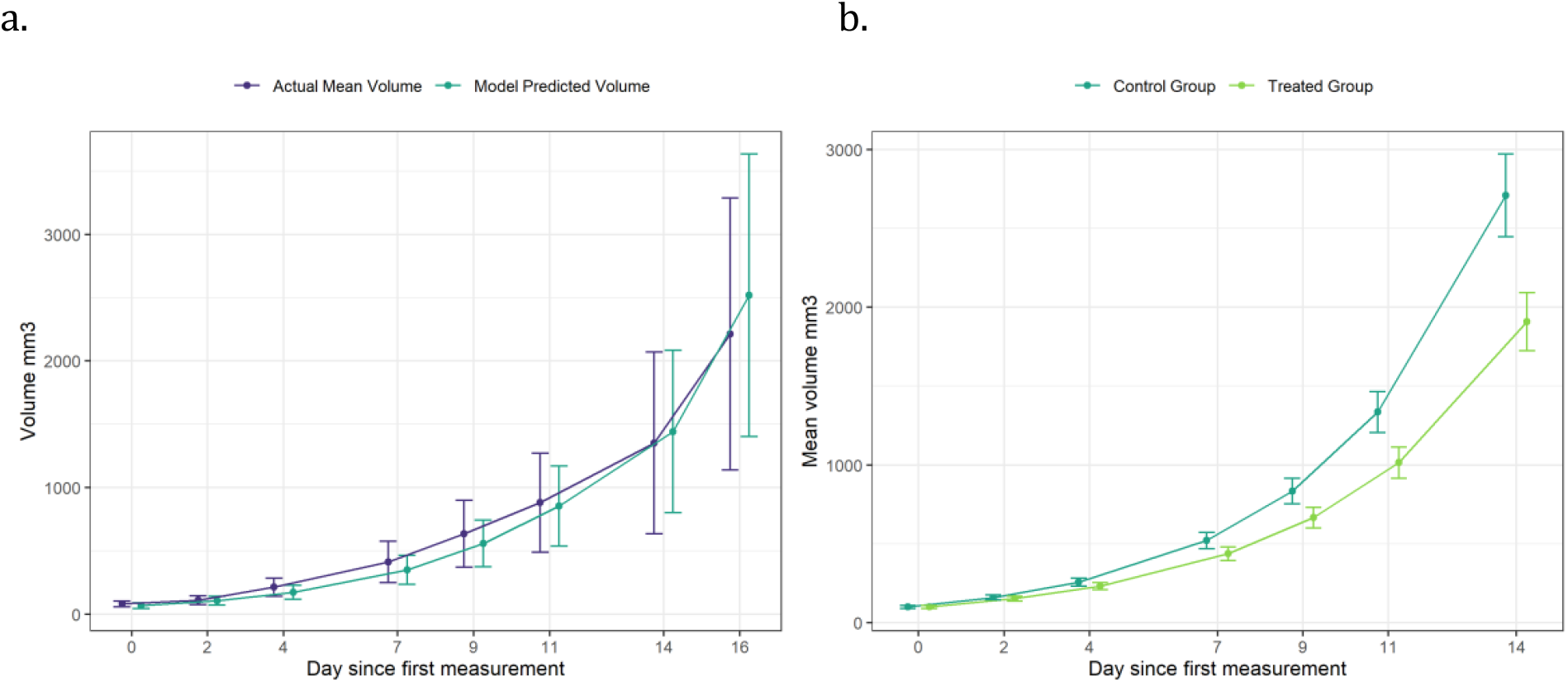
**a. Comparison of study average tumor volume against model predicted average volumes.** Model used to predict rodent volumes averaged across the 5 users. Error bars are 95% CIs **Figure 2. b. Average tumor growth of representative synthetic study data**. Using the growth rate obtained from the model, synthetic study data was created in which a treatment was evaluated and initial group volumes across the groups were equal. This is referred to as the “ground truth study” and was the basis for which the impact of user variability was investigated. Error bars are 95% CIs

Using the growth rate obtained from the model, synthetic study data were created. Synthetic study data with unwanted sources of variability removed was essential to be confident in the conclusions made. The data were created with the following changes to the original study:

- A treated group was added which the same growth rate as the control group minus a set value. This value was then varied to adjust the growth rate, creating different treatment effectiveness levels.
- 8 rodents were created in each group and tumor volumes were initially identical across the two groups. This was to replicate the effects of randomization where initial group volumes are equal and any difference in group volume after treatment can be attributed to treatment alone. Initial rodent tumor volumes are shown in **Table 1**.

**Table 1:** Initial rodent volumes for both groups in the synthetic study

| *Rodent Name* | 01 | 02 | 03 | 04 | 05 | 06 | 07 | 08 |
| --- | --- | --- | --- | --- | --- | --- | --- | --- |
| *Initial Tumor Volume (mm<sup>3</sup>)* | 80 | 85.7 | 91.4 | 97.1 | 102.9 | 108.6 | 114.3 | 120 |

After establishing the initial conditions, the model was used to simulate the growth of these rodents in both groups for the 7 measurement sessions across 14 days (**Figure 2.b**). These synthetic study data will be referred to as the ground truth study.

### Creating different treatment scenarios

To investigate the impact of inter-operator variability at different treatment levels, three separate ground truth studies were created, each with a different growth rate in the treated group to represent different treatment strengths or dosages. The treated group growth rate was defined as a constant and subtracted from the control group growth rate:

- Most effective treatment scenario = 0.236 – 0.03
- Effective treatment = 0.236 – 0.025
- Least effective = 0.236 – 0.02

On the final day this resulted in the following group mean difference and standard errors:

- Most effective treatment scenario: Mean group difference = 950mm^3^
- Effective treatment: Mean group difference = 800mm^3^
- Least effective: Mean group difference = 650mm^3^

Both the control and treated groups in each scenario had a standard error ∼ 120mm^3^.

A t-test was performed on the final day to determine if there was a significant difference between average group volumes in the ground truth study. In all 3 of these scenarios (where unwanted variability was equal to zero), the treatment group was statistically different to the control group (**Table 2**). To investigate the impact of inter-operator variability on the outcome of a study, user variability was then introduced to the ground truth study data to determine how this variability affected the ability to statistically separate the two groups.

**Table 2:** Results of t-test comparing final day group volumes in each of the three ground truth studies

| *Treatment Scenario* | *p-value* |
| --- | --- |
| Most Effective | $8 \times 10^{-5}$ |
| Effective | $1 \times 10^{-4}$ |
| Least Effective | $3 \times 10^{-4}$ |

## Defining user variability

### In vivo data processing

Percentage difference from the mean (PDFM) was used to quantify how users measure in relation to each other. The PDFM was modified to exclude a user’s own measurements, so that users were not compared to themselves. PDFM is a relative measure affected by tumor size so to accurately simulate user variation throughout a longitudinal study, PDFM was assessed for a range of tumor sizes. Therefore, only longitudinal studies that captured the entire tumor life cycle (from palpable to welfare endpoint) and that had at least 3 users taking repeat measurement were included. 9 studies in our global dataset met these criteria, summarized in **Table 3**.

**Table 3:** Study data used to compute PDFM and users which measured in the studies

| *Study* | *Users* |
| --- | --- |
| Study 1 | u01, u02, u03 |
| Study 2 | u01, u02, u06 |
| Study 3 | u04, u05, u07 |
| Study 4 | u08, u09, u10, u11, u12 |
| Study 5 | u08, u09, u10, u11, u12 |
| Study 6 | u08, u09, u10, u11, u12 |
| Study 7 | u13, u14, u15 |
| Study 8 | u13, u14, u15 |
| Study 9 | u13, u14, u15 |

PDFM was calculated for each user with each measurement device and across three size ranges:

- Small tumors <= 200mm^3^
- Medium tumors: between 200 and 800mm^3^
- Large Tumors: >= 800mm^3^

These bounds were chosen to maximize the number of datapoints across the size ranges whilst still being somewhat representative. **Table 4** details the number of data points (unique measurements) included in each size range.

**Table 4:** Number of data points for the various size ranges for both 3D-TI and Calipers. T.V = Tumor Volume

| *Measurement Device* | *Tumor Size Range* | *Number of Data Points* |
| --- | --- | --- |
| 3D-TI | Small: $\text{T.V} \leq 200\text{mm}^3$ | 1028 |
| 3D-TI | Medium: $200\text{mm}^3 < \text{T.V} < 800\text{mm}^3$ | 1073 |
| 3D-TI | Large: $\text{T.V} \geq 800\text{mm}^3$ | 1362 |
| Calipers | Small: $\text{T.V} \leq 200\text{mm}^3$ | 1177 |
| Calipers | Medium: $200\text{mm}^3 < \text{T.V} < 800\text{mm}^3$ | 1278 |
| Calipers | Large: $\text{T.V} \geq 800\text{mm}^3$ | 1001 |

#### Capturing characteristics of users for 3D-TI and Calipers

PDFM was used to establish whether a user had a particular bias when compared to other users, e.g., User 03 had a mean PDFM of ∼50% for large tumors with calipers and as such typically measured 50% larger than the average of the other users’ measures in the same study (**Figure 3a**). Consistency of said biases relative to the other users in the same study is also an important factor to consider and was assessed using the standard deviation of PDFM. 3D-TI users were less biased than caliper users for all size ranges when assessed by the mean (**Figure 3.b**). Users were also more consistent in their biases when using 3D-TI as opposed to calipers when looking at the standard deviation for large and medium size tumors (**Figure 3b**, p = 8.3 × 10^−5^ and p = 1.6 × 10^−2^ respectively, t-test). **Figure 4** shows examples of user measurement bias as well as inconsistency of bias.

**Figure 3.**
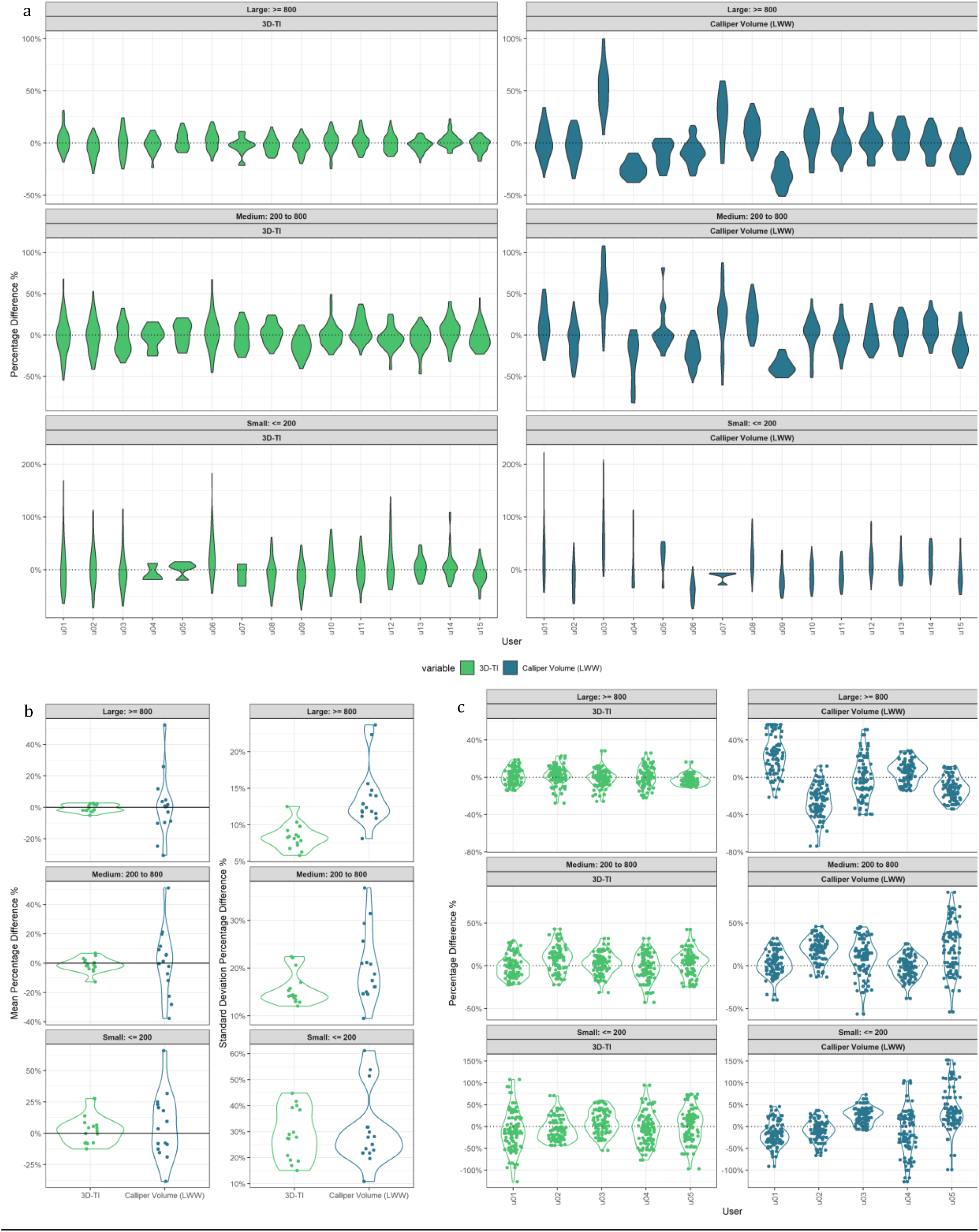
**a. Percentage difference from the mean (PDFM) for all users.** Shown as a violin plot, users taken from valid longitudinal studies for 3D-TI (Left) and Calipers (right) for a range of tumors sizes. **Figure 3.b. Mean (left) and standard deviation (right) of PDFM of each user**. Violin plot shown to emphasize distributions, split by tumor size range **Figure 3.c. Generated PDFMs for 5 synthetic users for 3D-TI (left, green) and Calipers (right, blue)**. To create synthetic user data for a given size range, randomly sample a relevant mean and standard deviation from 3.b. Use this in a normal distribution to generate 50 new PDFMs.

**Figure 4:**
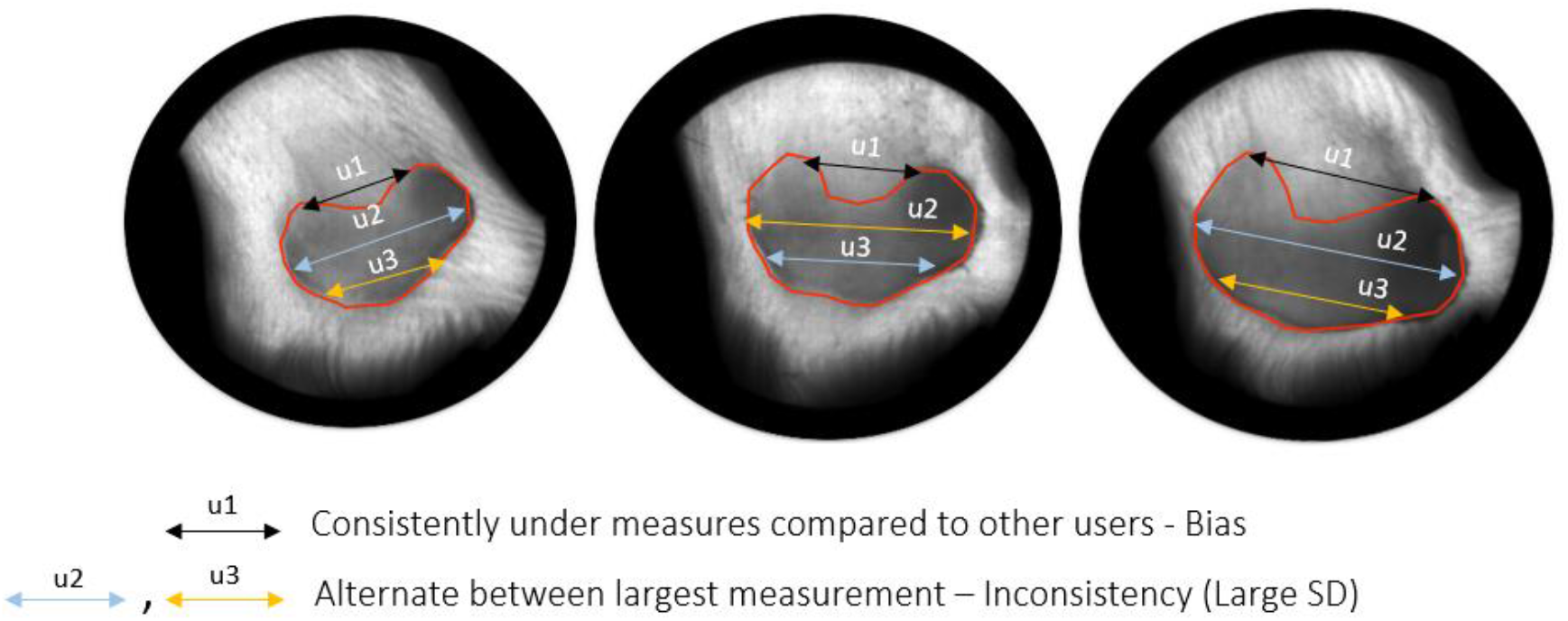
Diagram illustrating user bias (user 1) and inconsistency of bias compared with other users (users 2 and 3).

#### Generating synthetic users

100 synthetic users were generated from measurement data of 15 original users to produce more robust results and reduce effects of outliers (**Figure 1, box 1b**). To generate each synthetic user, a mean and standard deviation from **Figure 3.b** was randomly sampled from each size range for each device. The mean and standard deviation were then used in a normal distribution to generate 50 percentage differences for that synthetic user.

Over half the percentage difference distributions for each user, size range and device (3D-TI and calipers) were not normally distributed when assessed using a Shapiro-Wilk test due to large outliers. Therefore, the top and bottom 5% of percentage differences were trimmed for each user, device and size range to exclude outliers and reduce the number of not-normal cases to 1/3. **Figure 3.c** shows generated percentage differences for 5 example synthetic users. We can see that they share similar trends to the percentage difference plots shown in **Figure 5.a**.

**Figure 5:**
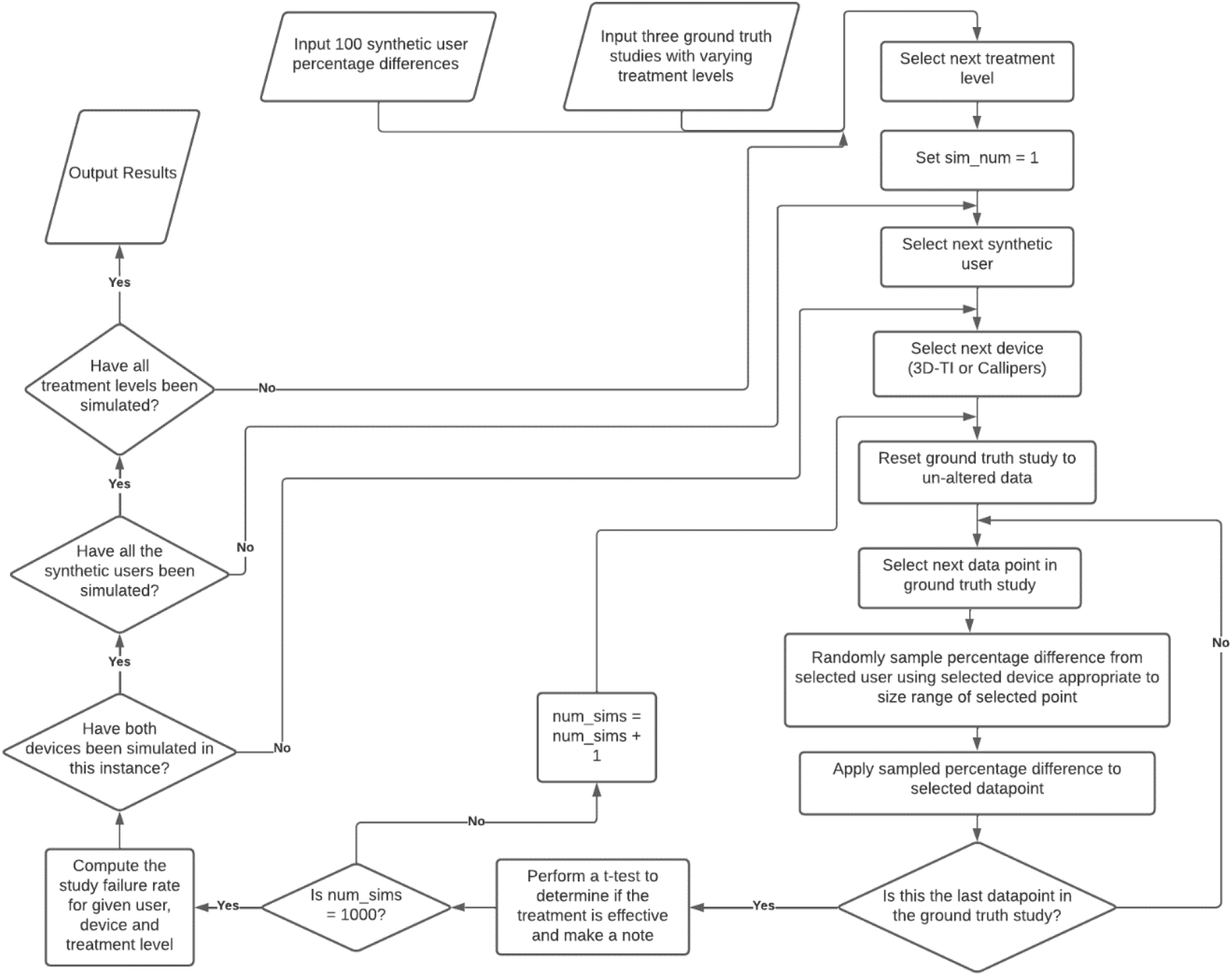
Flow chart detailing the simulation algorithm

#### Simulating user measurement in the ground truth study

After generating both the ground truth study data and 100 synthetic users that accurately represented 3D-TI and Caliper users, the two were then combined to investigate the impact of inter-operator variability on study outcome (**Figure 1, box 2**).

The ground truth study data were categorized according to tumor volume using the same ranges (**Figure 3**). An iterative process was then run in which 1 of the 100 synthetic users was selected and their percentage differences were applied to the ground truth study data at the appropriate size range. This was simply done by selecting a datapoint in the ground truth study, randomly sampling one of the percentage differences within the same tumor size category and multiplying them together, essentially “reverting” back from a mean measurement as generated by the model to the individual user measurement. This was repeated for every data point in the ground truth study, for both the generated 3D-TI and caliper percentage differences. The synthetic study data was then determined to have been “measured” by the selected user. A t-test was then performed on the final day to determine if there was a significant difference in average volume between both groups. This process of synthetic measurement and analysis was repeated for each user 1000 times (for 3D-TI and caliper measurements) to sample widely and create a stable mean. An incorrect result rate was computed as the number of times the control and treatment groups could not be statistically separated divided by 1000 for each user and each device.

The above simulation was performed for each of the three treatment scenarios. The study failure rate was then averaged across all 100 users, split across 3D-TI, calipers, and the three treatment levels (**Figure 5**).

## Results

An existing longitudinal in vivo study was used to create synthetic tumor growth curve data for a range of treatment scenarios as detailed in Methods. Inter-operator measurement variability was then computed for both 3D-TI and calipers and applied to the synthetic growth curves to generate user measurements in silico. These data were then analyzed in order to investigate the effect of user variability on study endpoint.

For a range of treatment scenarios, 3D-TI consistently reduced the chance of getting an incorrect result in an efficacy study (**Table 5**). Failing to detect a significant difference between group means on the final study day was classed as an incorrect result (false negative). For the most effective scenario where the difference in mean group volumes between control and treated groups was 950mm^3^ before applying user variability, the treatment was deemed effective for all 100 simulated 3D-TI users and their 1000 measurement repeats. When using calipers in the same treatment scenario, an incorrect result (false negative) was obtained 3.5% of the time. When decreasing the effectiveness of the treatment, inter-operator variability had an increasing impact on endpoint results. The chance of getting an incorrect result was more than doubled for calipers in the “effective” scenario at 7.6%. For 3D-TI measurements, an incorrect result was only obtained in 0.2% of cases. Finally for the scenario in which the treatment is the least effective of the three, caliper measurements obtained an incorrect result in almost 20% of cases. For 3D-TI, even in the least effective treatment scenario an incorrect result was only obtained in 2.4% of cases.

**Table 5:**
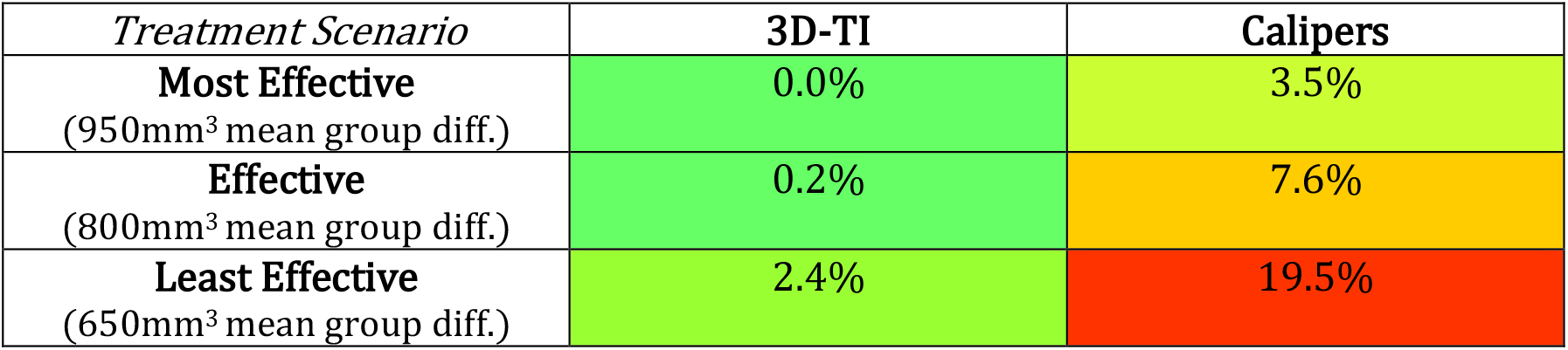
Probability of incorrectly determining an effective treatment to be ineffective due to user variability for 3D-TI and Calipers. Using a combination of simulation and modelling, 100 synthetic users “measured” on a study using both 3D-TI and Calipers in which the treatment was known to be effective, this process was repeated 1000 times for each user. Probability of not detecting a difference between group means was computed for each user then averaged across all users for a given device. Volumes in brackets is the mean group difference for the respective treatment scenario.

| *Treatment Scenario* | **3D-TI** | **Calipers** |
| --- | --- | --- |
| **Most Effective**<br>(950mm<sup>3</sup> mean group diff.) | 0.0% | 3.5% |
| **Effective**<br>(800mm<sup>3</sup> mean group diff.) | 0.2% | 7.6% |
| **Least Effective**<br>(650mm<sup>3</sup> mean group diff.) | 2.4% | 19.5% |

Figure 6. highlights the difference in characteristics between the 5 users with the lowest incorrect result rate and the 5 highest users. Users that measured less consistently when compared with other users and therefore had larger standard deviation in PDFM, were more likely to obtain an incorrect result. Users with mean PDFMs centered about zero were also likely to produce an incorrect result as both positive and negative PDFMs created more overlaps in the group measurements.

**Figure 6:**
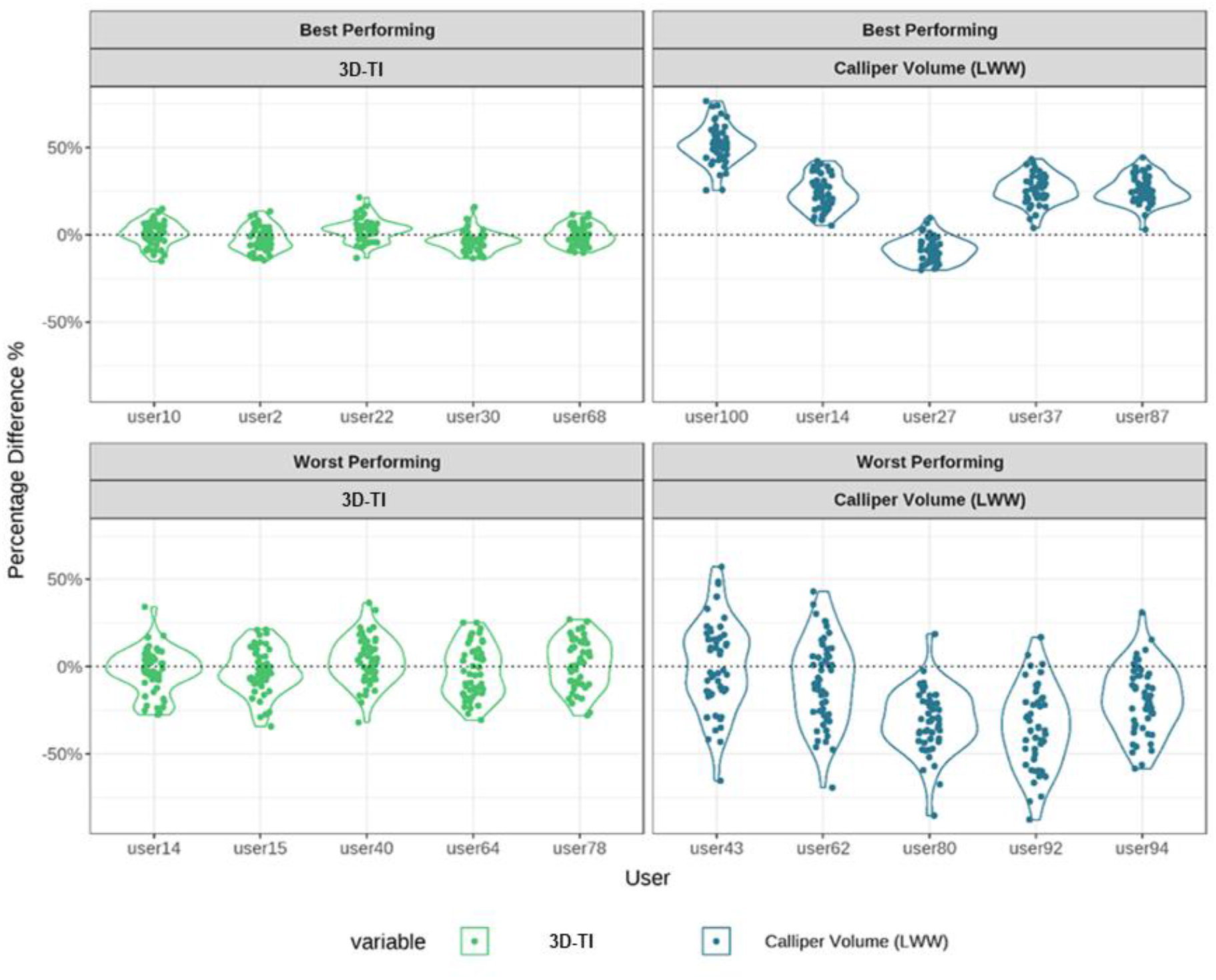
Percentage difference from the mean (PDFM) for large tumors. Data plotted as violin plots for the 5 synthetic users with the lowest (top) and highest (bottom) incorrect result rates for 3D-TI (left) and Calipers (right).

The impact of mean PDFM and the standard deviation of PDFM on incorrect result rate was then investigated further, taking into account all 100 synthetic users, for 3D-TI and Calipers (**Figure 7**). Users with larger standard deviations had a higher chance of obtaining an incorrect result, and caliper users tended to have larger standard deviations. As the standard deviation of PDFM increased, users who did not over measure were more likely to obtain an incorrect result in this scenario. The best user is one who has little to no bias (mean of 0) and is highly consistent compared to other users (small standard deviation).

**Figure 7:**
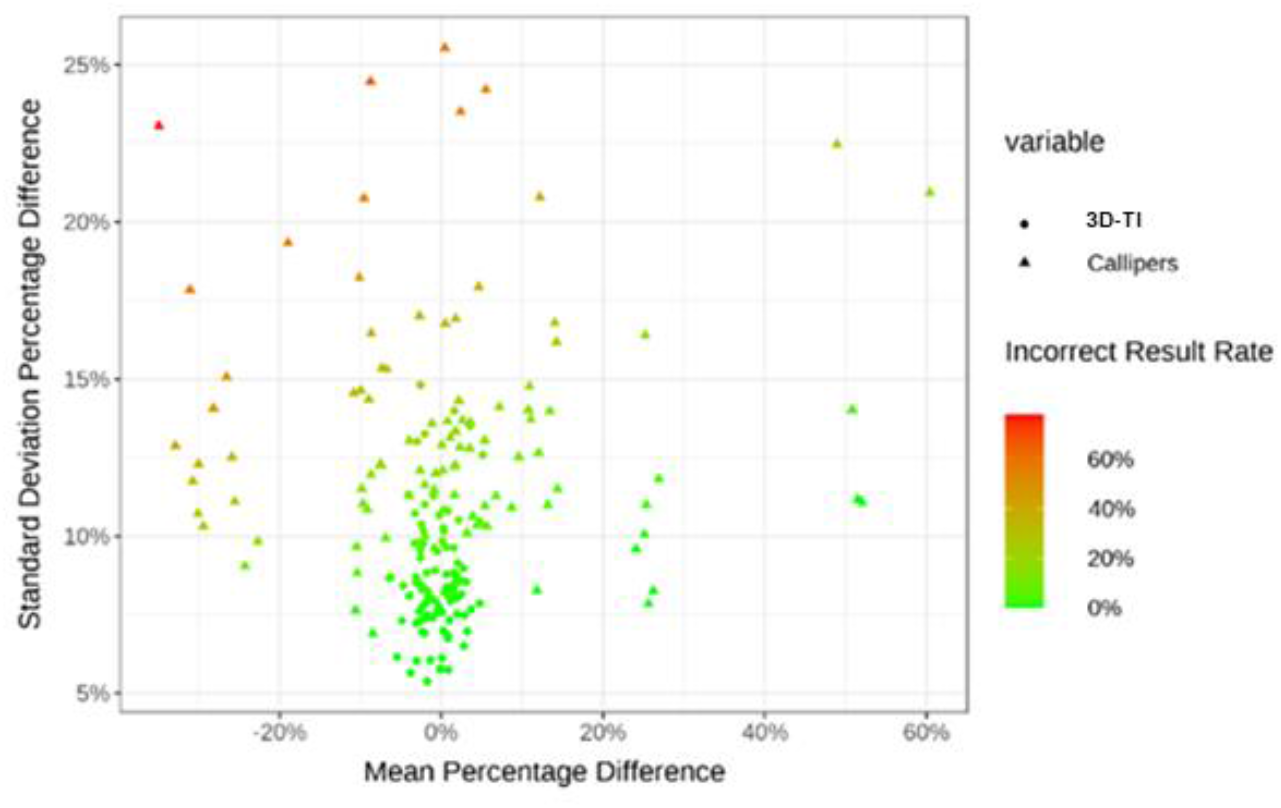
Standard deviation vs mean of PDFM. Mean and standard deviation PDFM colored by incorrect result rate for all 100 synthetic users using 3D-TI and Calipers. Data for large tumors only.

## Discussion

Preclinical study irreproducibility stems from many causes, of which user measurement variability has been shown to be a significant problem when taking tumor measurements using calipers^2–5^. Gathering repeat measurements (either inter-or intra-operator) is time consuming and increases welfare concerns due to increased animal handling, so for this study, real in vivo data was used to create a synthetic model. Working in silico also allowed exclusion of all other sources of in vivo variability such as differences in group means at randomization. Thus, user variability was isolated and was the sole cause of any change in study outcome.

The synthetic user section of the analysis was taken from real data obtained from 9 longitudinal studies, capturing a wide range of tumor sizes. Our model assumes that the means and standard deviations of each user’s percentage differences can be interchanged to form new users. It may be that particular cell lines result in a larger standard deviation than others and shouldn’t be compared/interchanged with characteristics from other cell lines. To further improve the model in future, it may be useful to better define how the mean and standard deviation are affected by other variables including cell line.

PDFM outliers were excluded from the model to better generalize the PDFM as a normal distribution, however this method slightly underestimates the true impact of user variability on study outcome. We predict that including outliers would further increase the chance of false results from the values reported here. On the other hand, sampling percentage difference randomly from one synthetic user, and resampling in a study repeat may overestimate variation in comparison to real-life measurement; it is unlikely that one user would repeat a measurement and record a greatly different result when resampling.

Interestingly, a user who over measures relative to other users is less at risk at detecting a false negative than a user which under measures or measures consistently with other users, as shown by the effect of mean PDFM on the incorrect result rate. The reason for this is large positive percentage differences creates a larger distance in group means than negative similar sized percentage differences. The inverse would be true with a scenario with a treatment that was not effective and false positive rate were to be investigated.

This study is a starting point for further investigations into the effects of measurement variability on study endpoints and outcomes. Greater understanding of these effects will help us to understand and minimize problems in study reproducibility and to accurately characterize drug effects at an early stage with confidence. We have shown here that reducing measurement variability reduces false negative rate which has been identified as an essential variable to control to achieve cost-savings in clinical trials using the ‘quick win, fast fail’ model^19^, without mistakenly excluding effective drugs from further development.

Next, investigating the impact of inter-operator variability study on false positive rate should be reported to get a full understanding of overall false result rates. Secondly, a wider range of different treatment evaluation scenarios could be investigated, including regression studies, and impact on other efficacy measures such as survival curves, and other statistical tests such as a repeated measures ANOVA. More in-depth analysis to determine the effects of cell line on inter-operator variability would also enhance understanding of specific treatment scenarios.

Calipers are by far the cheapest tumor measurement tool used universally by researchers and animal technicians working in the oncology field. Hesitancy to change a long-established and ubiquitous technique, as well as additional welfare considerations (anesthesia requirement) are barriers to wider adoption of alternative imaging techniques. High start-up costs are another factor, but one that could be offset over time by new technologies that increase throughput by automating data collection and entry, and which have the potential to reduce group sizes and study length by offering more accurate and precise data.

In conclusion we showed that by using a 3D and thermal imaging device to reduce user measurement variability in comparison to calipers, the chance of a false efficacy study result was also decreased. This translates to missing a treatment effect in an efficacy study, and wrongfully excluding a viable drug candidate from further development. The inverse is also possible; a false positive result also has the potential to be a costly mistake if an ineffective drug moves forward in the development cycle for more rounds of testing. Resources and time would be wasted trying and eventually failing to replicate the false positive result. Later-stage clinical trials are expensive and time consuming to run, therefore incorrect or ambiguous results should be reduced as early on in development as possible.

## Acknowledgements

The authors thank and gratefully acknowledge the contribution of all the scientists who ran studies and collected the measurement data which was analyzed in this report. We also thank Adam Sardar for his guidance and discussion of the modelling methods.

